# Correcting for Body Size Variation in Morphometric Analysis

**DOI:** 10.1101/2021.05.17.444580

**Authors:** Kin Onn Chan, L. Lee Grismer

**Affiliations:** Lee Kong Chian Natural History Museum, 2 Conservatory Drive, 117377 Singapore; Herpetology Laboratory, Department of Biology, La Sierra University, 4500 Riverwalk Parkway, Riverside, California 92505, USA

**Keywords:** Allometry, Allometric growth, Body size, Continuous trait, GroupStruct, Morphology, Principal components analysis, Size correction

## Abstract

Using an allometric growth model to correct for body size variation has been known for many decades to be superior to several other widely used methods such as ratios, analysis of covariance, principal components analysis, and residual analysis. However, this technique remains relatively obscure and rarely applied. We optimize the implementation of this method through a newly developed and easy-to-use R package *GroupStruct* and use empirical datasets to test its relative efficacy compared to several commonly used methods. Our results demonstrate the superiority of the allometric method and highlights the negative impacts of applying improper body size correction methods.

## INTRODUCTION

Most morphological traits scale with body size, hence, observed differences in morphological characters are intrinsically associated with differences in overall body size. Variation in body size and shape can be influenced by many factors including but not limited to ontogeny (growth), geography, sexual dimorphism, environment, and evolutionary history (Stillwell, Morse, & Fox, 2007; Calderón-Espinosa, Ortega-León, & Zamora-Abrego, 2013; Grismer & Grismer, 2017; Chan *et al.*, 2018; Chole, Woodard, & Bloch, 2019). Conflating multiple axes of variation can therefore have confounding effects that can bias results. For example, in studies where morphometric data are used to distinguish between species, it is crucial to correct for ontogenetic variation to ensure that differences in morphological characters reflect distinct evolutionary histories as opposed to being conflated with variation in body size. This is an important step because most empirical morphometric datasets contain ontogenetic heterogeneity that is unavoidable when sampling natural populations or archival material.

Among the most widely implemented methods to correct for ontogenetic size variation are ratios, *x/y*, where *x* is the dependent character and *y* is the standard measure of growth stage such as body length or snout-vent-length (Chan, Grismer, & Brown, 2014); residual analysis, where the dependent character (*x*) is size-corrected according to the standard measure (*y*) by obtaining the residuals from a linear or least squares regression of *x* over *y* (Revell, 2009; Mahler *et al.*, 2010; Blackburn *et al.*, 2013); ANCOVA, where body size is specified as a covariate (Garcia-Berthou, 2001; McCoy *et al.*, 2006; Chan *et al.*, 2014); and principal components analysis (PCA), where the first principal component is assumed to be the latent size variable, while other axes are assumed to be independent of size (McCoy *et al.*, 2006). However, the use of these techniques are flawed due to the violation of assumptions. Ratios can only be used if growth is isometric, i.e. the relationship between the dependent and independent character is linear and passes through the origin. In other words, all body parts grow at the same rate, hence shape does not change with size (Thorpe, 1983; Lleonart, Salat, & Torres, 2000; Nakagawa *et al.*, 2017). Residual analysis assumes that growth is associated with a single independent character and confounds within- and between-group variation when independent variables are correlated (Thorpe, 1983; Garcia-Berthou, 2001; Freckleton, 2002; McCoy *et al.*, 2006). PCA-based correction suffers from statistical artifacts when the number of traits is few and the magnitude of variance is heterogeneous among traits (Thorpe, 1983; Berner, 2011). For more thorough reviews on these and other methods, see (Thorpe, 1976, 1983; Reist, 1985; McCoy *et al.*, 2006; Berner, 2011).

To address these shortcomings, a normalization equation that scales data according to an allometric growth model was outlined by Thorpe (1975) and further discussed by Reist (1985) and Lleonart *et al.* (2000). Although this method has been shown to be superior to other techniques (Reist, 1985), its usage has been relatively limited, presumably due to a more complicated procedure. As such, we optimize the implementation of this method through a newly developed R package *GroupStruct* (available at https://github.com/chankinonn/GroupStruct) and provide comparative and validation tests using empirical datasets.

## MATERIAL AND METHODS

The allometric formula outlined by Thorpe (1975) is as follows:

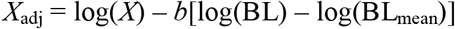

where *X*_adj_ = size corrected variable; *X* = unadjusted dependent variable; *b* = regression coefficient or slope of the relationship between log(*X*) and log(BL); BL = standard measurement of body length/size (e.g. snout-vent-length); BL_mean_ = grand mean of BL across species or populations. All logarithmic transformations are performed at base 10. This equation removes all information related to size, scales individuals to the same size, and adjusts their shape to the new size (Thorpe, 1983; Lleonart *et al.*, 2000). This adjustment is implemented through the *allom()* function in *GroupStruct*, which can handle multispecies and multipopulation datasets.

For *b*, two types of slopes can be estimated according to group structure: pooled groups combine individuals from different localities to produce a single compound locality, whereas common within-group pools individuals from the same population/locality (each population represents a separate pool). The operational taxonomic unit (OTU) in a multispecies dataset is represented by at least two species and multiple populations/localities of the same species are combined under a common OTU identifier. A slope is calculated for each species (pooled groups) and BL_mean_ represents the grand mean of each unique species. A multipopulation dataset comprises a single species where each unique OTU represents a different population. A slope is calculated for each population and BL_mean_ is the grand mean averaged across all populations (one single mean for the whole dataset).

We employed two empirical datasets (obvious juveniles removed) to demonstrate the use of *GroupStruct* and compare the efficacy of several widely used size correction techniques and their effects on evincing group structure. The first is a multispecies dataset comprising four species of Bent-toed geckos from the genus *Cyrtodactylus* from Thailand and Myanmar (*C. chaunghanakwaensis, C. taungwineensis, C. sinyineensis*, and *C. maelanoi*), all of which have been shown to be distinct species based on molecular and morphological data (Grismer *et al.*, 2018b,a, 2020b,a). A total of 13 mensural characters were assessed: snout-vent-length (SVL), pelvic width (PW), pelvic height (PH), axila-groin-length (AXL), head length (HL), head width (HW), head depth (HD), snout length (SNT), eye diameter (ED), hindlimb width (HDW), hindlimb length (HDL), forelimb width (FLW), and forelimb length (FLL). The second is a multipopulation dataset comprising three populations of *C. aequalis* from Myanmar that exhibit geographic variation (Grismer *et al.*, 2020c). This dataset contains nine mensural characters: SVL, FLL, AXL, HL, HW, HD, TBL (tibia length), EE (ear-to-eye distance), and ES (ear-to-snout distance). Sexual dimorphism was insignificant in both datasets (see references above), hence both male and female measurements were combined to boost sample size. In all analyses, the standard body measurement used to correct for body size was snout-vent-length (SVL). For regression analyses, SVL was set as the independent variable, while the other characters were set as dependent variables. To compare the performance of various body size correction techniques, we performed six different treatments on each dataset: (1) Raw: raw, unadjusted measurements; (2) Raw log: log-transformed raw measurements; (3) Ratio: each size variable was divided by SVL and subsequently log-transformed; (4) Residuals: residuals for each character were calculated using linear regression of each dependent variable against SVL; (5) PCA: principal components analysis was performed and the first principal component relating to size was discarded; (6) Allometry: characters were adjusted using the allometric equation implemented through the *allom()* function in *GroupStruct.* For the multipopulation dataset, we additionally analyzed data treatments with juveniles included to assess the effects of including juveniles in morphometric analyses. Here, we define juveniles as having SVLs that are less than half of the maximum SVL observed in a particular OTU.

The efficacy of various size correction methods was assessed by testing the correlation between the dependent character variables and SVL. Non-significant correlations indicate a successful removal of size effects. The impacts of size correction on multivariate analysis were examined using ANOVA and ANCOVA. We also performed principal components analysis (PCA) on each data treatment to investigate the performance of different size correction methods on evincing group structure. To quantify separation between groups in PC space and determine whether cluster separation is significant, we performed a Hotelling’s *T^2^* test on scores of the first two principal components (except the PCA dataset where the test is performed on PC2 and PC3). The *T^2^* statistic increases with increasing distance between two group centroids in PC space and with decreasing within-group variance. A significant *p*-value rejects the null hypothesis of no multivariate difference between groups, indicating a significant difference between PCA clusters. All analyses were performed in R (R Core Team, 2014). *GroupStruct* also contains the function *resid()* to calculate residuals and *ez_pca()* that performs PCA, summarizes the results, and produces PCA plots. The raw multispecies and multipopulation datasets and complete R script for all analyses performed in this study are available at https://doi.org/10.6084/m9.figshare.14605152.v1

## RESULTS

The correlation analysis shows that allometry is the most efficient at removing the effects of body size. After allometric adjustment on the multispecies dataset, the *p*-value is not significant (*p* = 0.08902) while the Multiple *R*^2^ and Adjusted *R*^2^ values are low, indicating a lack of correlation between the dependent character variables and SVL (Table 1). In contrast, *p*-values for the Ratios and Raw datasets remain significant (*p* = 0.016 and *p* = 0.000, respectively), indicating that the effect of size was is completely removed. For the multipopulation dataset, the Raw dataset remains highly significant after allometric adjustment (*p* = 0.000). Both Ratios and Allometry datasets have non-significant *p*-values *(p* = 0.1557 and *p* = 0.8587, respectively) but the significance and *R*^2^ values in the Allometry dataset are much lower compared to the Ratios dataset indicating a more efficacious removal of the size effect. Additionally, the relationships between each character and SVL in the Raw dataset are not linear and do not pass through the origin (Fig. 1), indicating the absence of isometry, thereby invalidating the use of ratios for size correction.

**Table 1.** Summary statistics of the correlation analysis of all dependent characters against SVL for the interspecific and intraspecific datasets. Non-significant *p*-values and low adjusted *R*^2^ indicate a lack of correlation between dependent characters and body size demonstrating the successful removal of size effects.

| | $F$-statistic | Multiple $R^2$ | Adjusted $R^2$ | $p$-value |
| --- | --- | --- | --- | --- |
| Mutispecies: |  |  |  |  |
| Raw | 106.1 | 0.9785 | 0.9693 | 0 |
| Ratios | 2.689 | 0.5049 | 0.3171 | 0.01621 |
| Allometry | 1.845 | 0.4415 | 0.2022 | 0.08902 |
| Multipopulation: |  |  |  |  |
| Raw | 78.25 | 0.9766 | 0.9641 | 0 |
| Ratios | 1.799 | 0.4896 | 0.2174 | 0.1557 |
| Allometry | 0.4552 | 0.1954 | -0.2338 | 0.8687 |

**Fig. 1.**
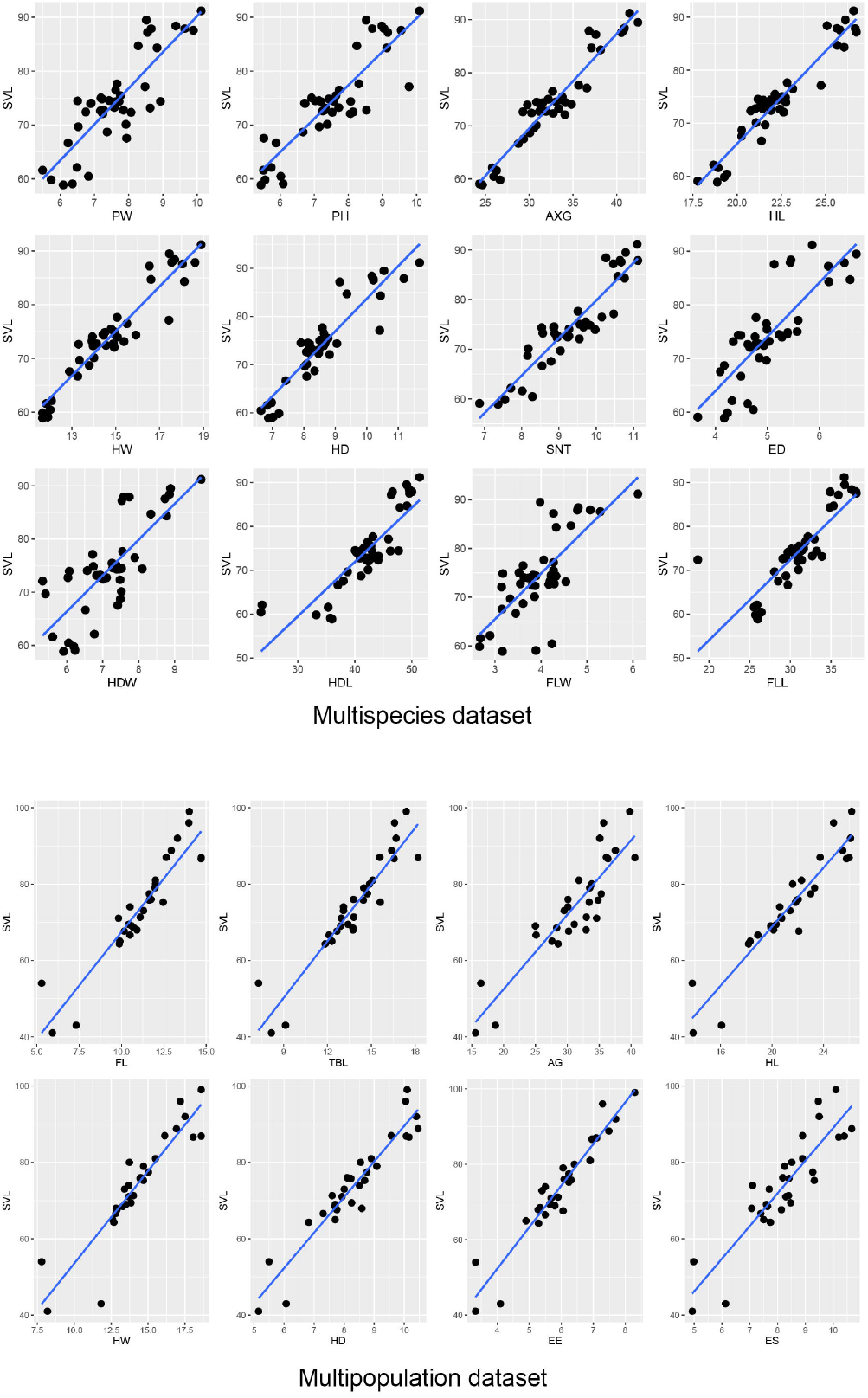
Scatter plots of each dependent character against SVL to test for isometric growth. The blue line represents the best-fit regression line. Deviation from the line and a slope intercept that does not pass through the origin indicates non-isometric growth.

The correlation analysis also demonstrates that all dependent characters are significantly correlated with each other, indicating that residuals should not be used to correct for body size (Tables S4, S5). The impact of different body-size correction methods on multivariate analyses is not inconsequential. ANCOVA on raw data and ANOVA on the ratios dataset produce relatively similar results with fewer variables that are significantly different (alpha = 0.001) compared to ANOVA on raw data (Fig. 2). However, ANOVA on the allometric dataset produces drastically different results where most characters are significantly different. Similarly, validation via PCA clearly shows that the allometric adjustment method performs the best in revealing group structure (Figs. 3–5) and produces the clearest separation across all datasets (Table 2). Log-transformation of raw measurements does not result in significant differences in the ordination of data points in PC space. The ratio and PCA methods for size removal perform worse in comparison to raw measurements as indicated by less separation of species clusters (Figs. 3–5). When juveniles are included, they form outliers and reduce the efficacy of all methods, including allometry (Fig. 5).

**Fig. 2.**
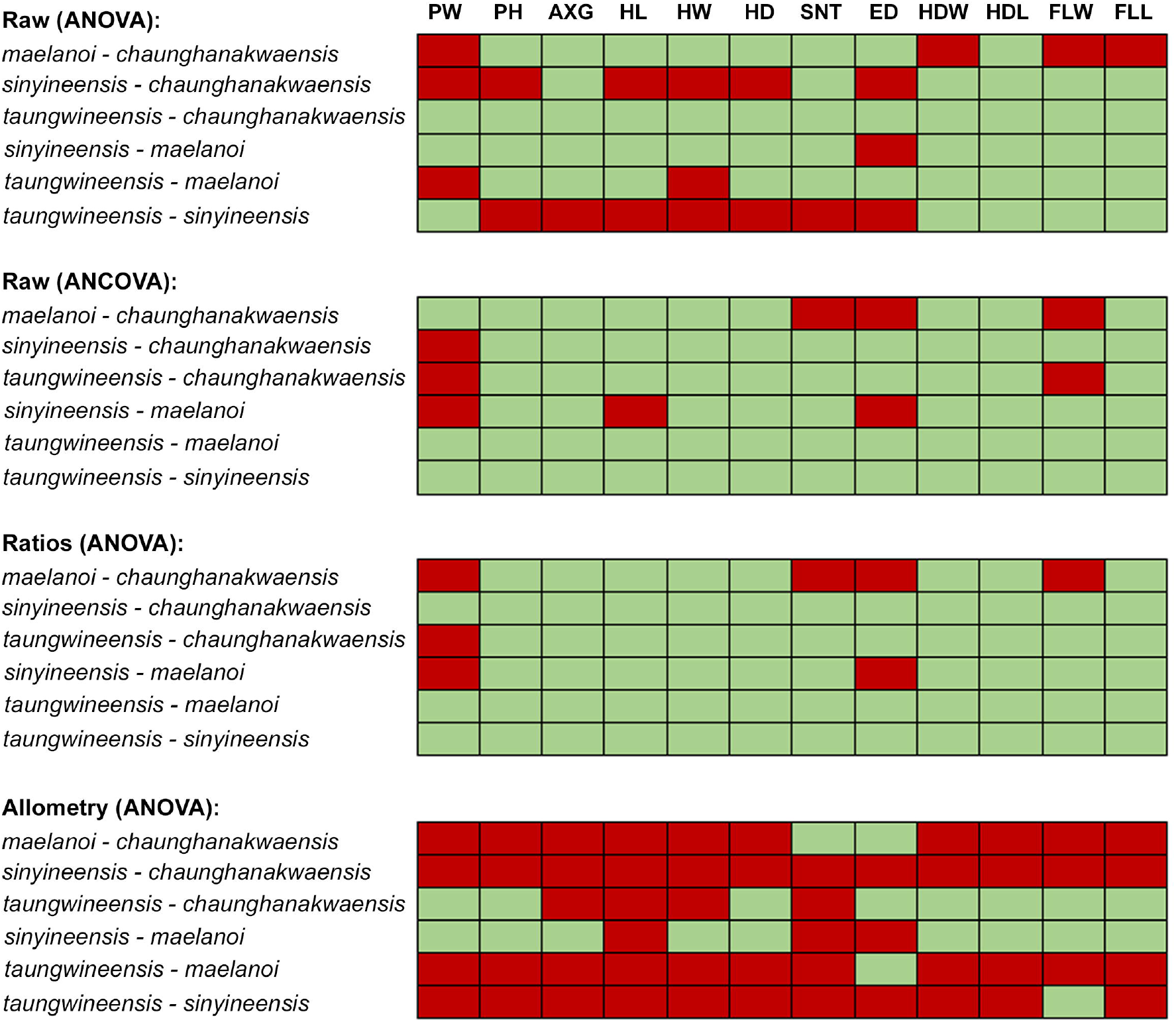
Results of ANOVA and ANCOVA performed on the raw, ratios, and allometric treatments on the interspecific dataset. Pairwise species comparisons for each dependent character were examined. Red boxes indicate *p* ≤ 0.001, green boxes represent *p* > 0.001.

**Fig. 3.**
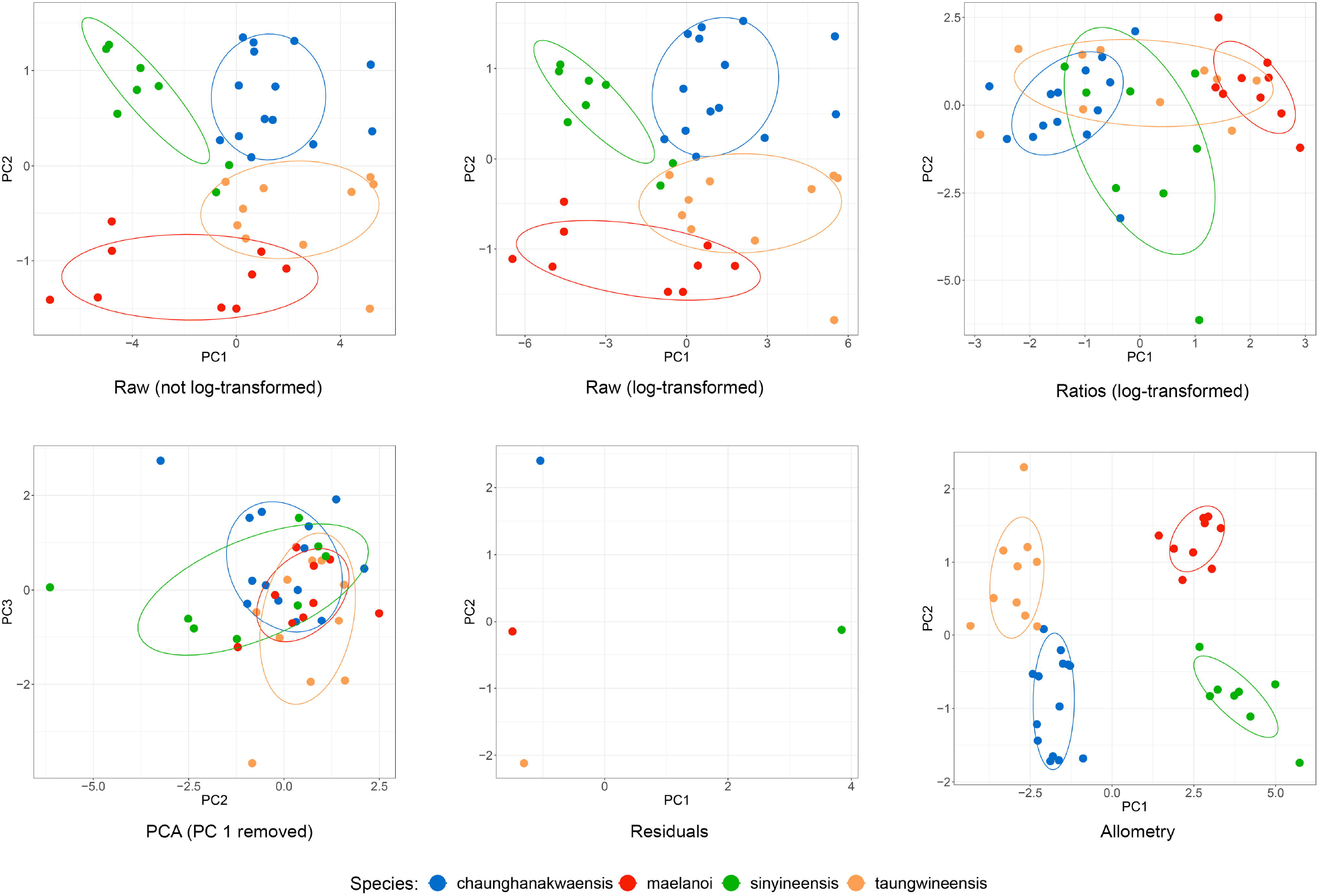
PCA plots of the various data treatments for the interspecific dataset that excludes juvenile measurements. All plots show PC1 against PC2 except for the PCA treatment, which plots PC2 against PC3 (PC1 was removed following the assumption that the first axis represents size).

**Fig. 4.**
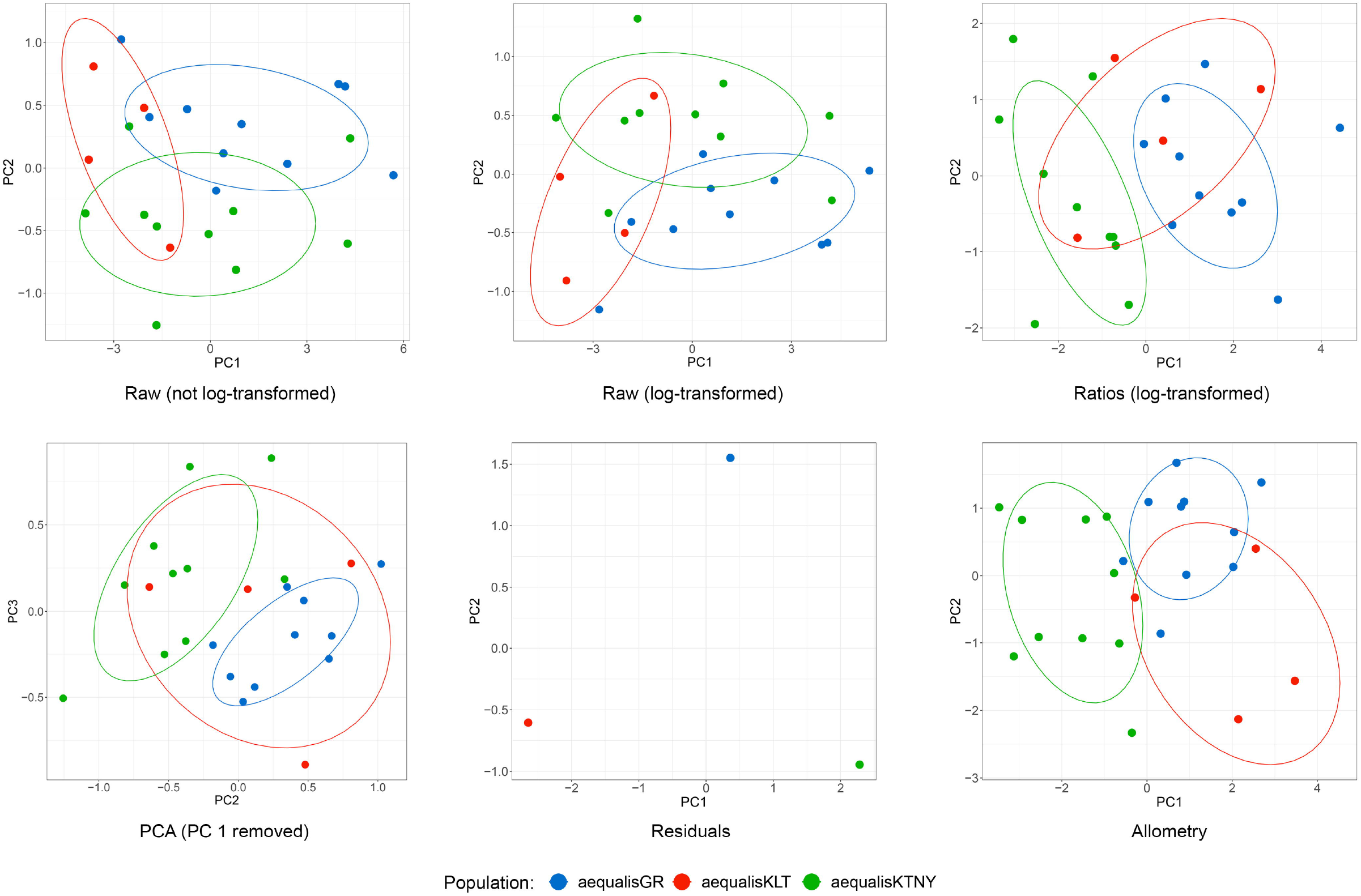
PCA plots of the various data treatments for the intraspecific dataset that excludes juvenile measurements. All plots show PC1 against PC2 except for the PCA treatment, which plots PC2 against PC3 (PC1 was removed following the assumption that the first axis represents size).

**Fig. 5.**
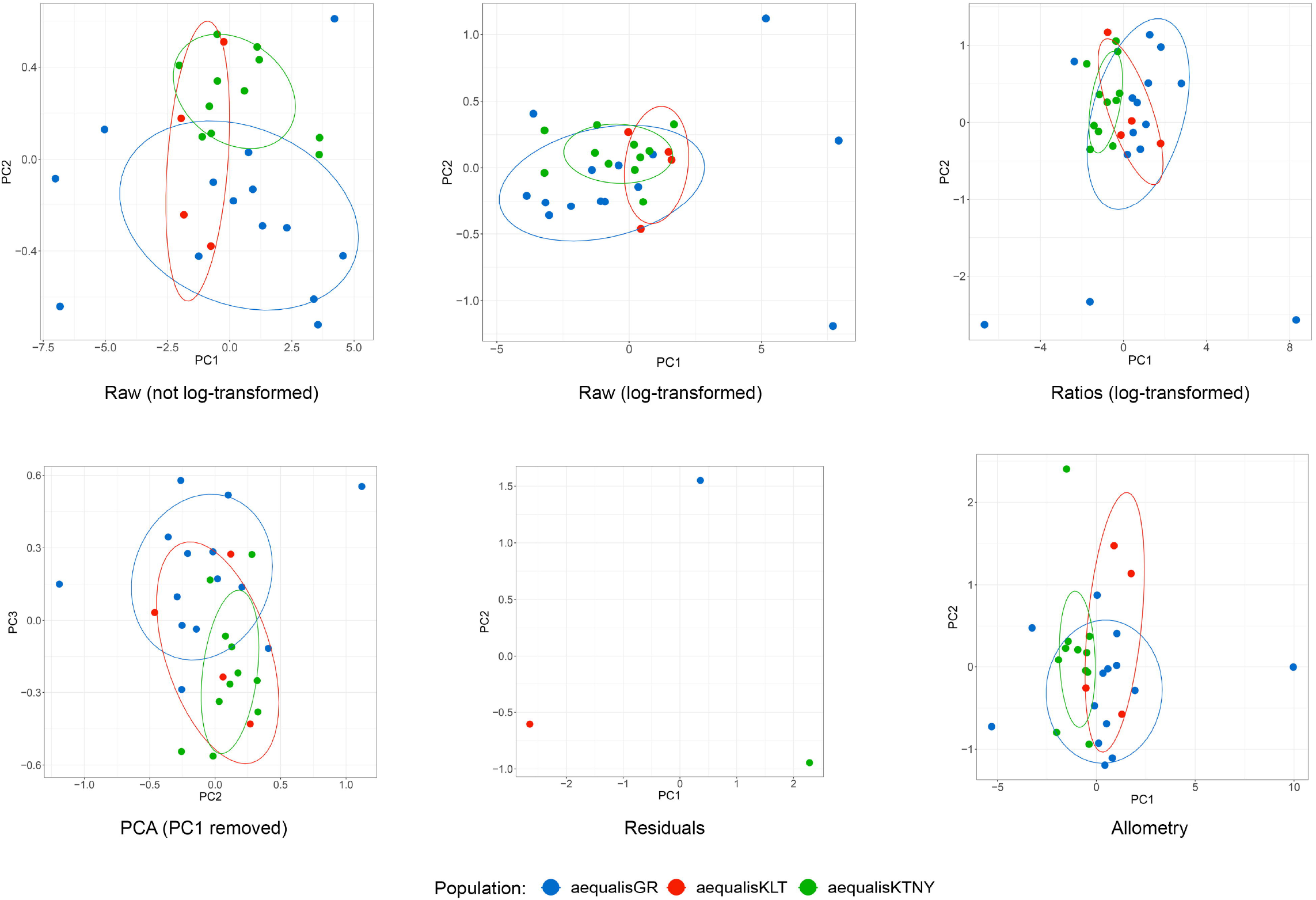
PCA plots of the various data treatments for the intraspecific dataset that includes juvenile measurements. All plots show PC1 against PC2 except for the PCA treatment, which plots PC2 against PC3 (PC1 was removed following the assumption that the first axis represents size).

**Table 2.**
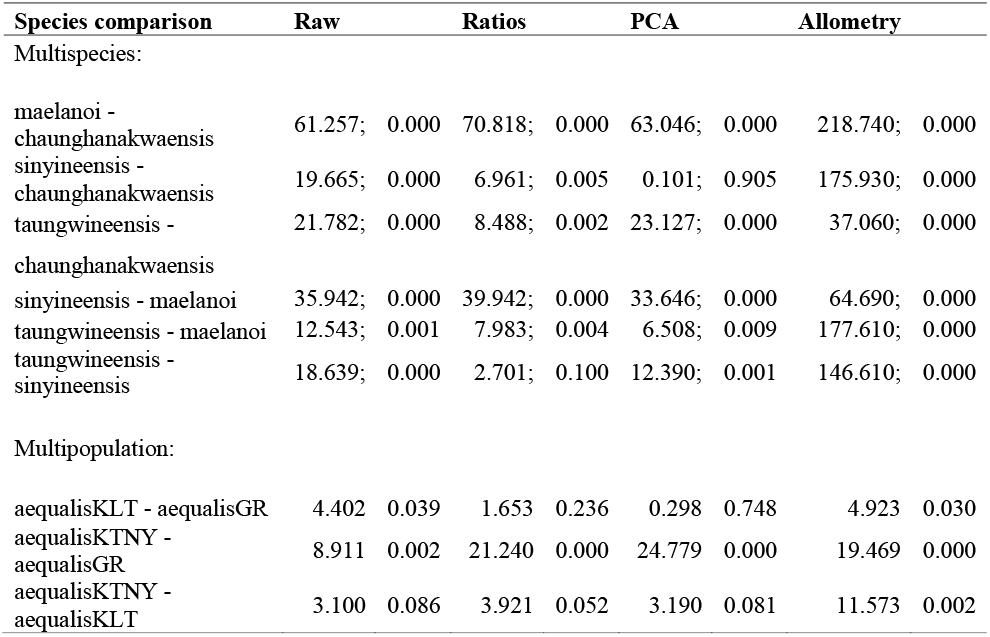
Results of the two-sample Hotelling’s *T^2^* test performed on the scores of the first two principal components (except the PCA dataset where the test was performed on PC2 and PC3) to quantify separation between groups in PC space. The *T^2^* statistic is followed by the *p*-value after the semicolon. The *T^2^* statistic increases with increasing distance between two group centroids in PC space and with decreasing within group variance. A significant *p*-value rejects the null hypothesis of no multivariate difference between groups, indicating a significant difference between PCA clusters.

| Species comparison | Raw | Ratios | PCA | Allometry |
| --- | --- | --- | --- | --- |
| Mutispecies: |  |  |  |  |
| maelanoi - chaunghanakwaensis | 61.257; 0.000 | 70.818; 0.000 | 63.046; 0.000 | 218.740; 0.000 |
| sinyineensis - chaunghanakwaensis | 19.665; 0.000 | 6.961; 0.005 | 0.101; 0.905 | 175.930; 0.000 |
| taungwineensis - | 21.782; 0.000 | 8.488; 0.002 | 23.127; 0.000 | 37.060; 0.000 |

|  |  |  |  |  |  |  |  |  |
| --- | --- | --- | --- | --- | --- | --- | --- | --- |
| chaunghanakwaensis |  |  |  |  |  |  |  |  |
| sinyineensis - maelanoi | 35.942; | 0.000 | 39.942; | 0.000 | 33.646; | 0.000 | 64.690; | 0.000 |
| taungwineensis - maelanoi | 12.543; | 0.001 | 7.983; | 0.004 | 6.508; | 0.009 | 177.610; | 0.000 |
| taungwineensis -<br>sinyineensis | 18.639; | 0.000 | 2.701; | 0.100 | 12.390; | 0.001 | 146.610; | 0.000 |

Multipopulation:
|  |  |  |  |  |  |  |  |  |
| --- | --- | --- | --- | --- | --- | --- | --- | --- |
| aequalisKLT - aequalisGR | 4.402 | 0.039 | 1.653 | 0.236 | 0.298 | 0.748 | 4.923 | 0.030 |
| aequalisKTNY -<br>aequalisGR | 8.911 | 0.002 | 21.240 | 0.000 | 24.779 | 0.000 | 19.469 | 0.000 |
| aequalisKTNY -<br>aequalisKLT | 3.100 | 0.086 | 3.921 | 0.052 | 3.190 | 0.081 | 11.573 | 0.002 |

## DISCUSSION

The importance of removing size effects from morphometric analyses cannot be understated and our results provide validation for the superiority of the allometric method over other widely-adopted techniques such as ratios, residuals, ANCOVA, and PCA. Our study demonstrates one example in which body size variation can produce spurious diagnoses of species boundaries. Allometric adjustment was able to reveal significantly more differences among characters (Fig. 2) and not only yielded better separation among species in the multispecies dataset but also revealed more nuanced attributes. For example, in the raw treatment, there was no separation along PC1, while the blue and green clusters were separated from the red and orange clusters along PC2 (Fig. 3). After allometric adjustment, orange and blue clusters were separated from red and green clusters along PC1, while orange and red clusters were separated from blue and green clusters along PC2, indicating that more variation was captured along both axes.

In our datasets, the assumptions of isometry and character independence were violated, which invalidated the use of ratios (Thorpe, 1983; Lleonart *et al.*, 2000; Nakagawa *et al.*, 2017) and residuals (Thorpe, 1983; Garcia-Berthou, 2001; Freckleton, 2002; McCoy *et al.*, 2006), respectively. Additionally, the residual analysis produces a single value per group, which may be appropriate for studies requiring a single point estimate per OTU (Revell, 2009; Mahler *et al.*, 2010; Blackburn *et al.*, 2013) but less suitable for studies focusing on the breadth of variation (e.g. diagnosing OTUs). We argue that many empirical datasets derived from natural populations violate the assumptions of isometry and character independence, which should be tested before ratios and residuals are used for size correction.

The inclusion of juveniles severely reduced the efficacy of all size correction methods including allometry, which is supposed to correct for ontogenetic variation (Fig. 5). This is because juvenile measurements can drastically skew the value of BL_mean_. Furthermore, juveniles appear to deviate more from isometric growth (Fig. 1). In other words, the large difference in juvenile morphology is treated as if data from two differently shaped species are conflated. Another crucial parameter in the allometric equation is the regression coefficient or slope. The accuracy of this parameter may not be severely affected by juvenile measurements but can be biased by a low sample size. Therefore, for the allometric method to be accurate, juveniles should be excluded and sample sizes for each OTU should be sufficiently large. Applying allometric size correction on datasets comprising samples with uncertain taxonomic rank (species vs. population) can also be an issue because the multispecies and multipopulation implementation of this method can produce different results. In such a case, we recommend that users apply the multispecies method that can better account for potential differences in average body size among OTUs.

Decoupling shape from size variation is desirable not just in the context of species diagnoses but for virtually every study that seeks to understand morphological variation among groups. This is particularly important in taxonomic studies where validation analyses such as PCA or multivariate tests are used to corroborate and diagnose species boundaries delimited in molecular phylogenies (Grismer *et al.*, 2020c); studies inferring morphological phylogenies using paleontological data (Parins-Fukuchi, 2018); or phylogenetic comparative methods that combine phylogenies with phenotypic data to understand evolutionary processes such as the mode and tempo of trait evolution (Mahler *et al.*, 2010). The improper use of such data or lack thereof can lead to erroneous taxonomic conclusions (Rogell, Dowling, & Husby, 2020).

## ACKNOWLEDGMENTS

We thank Alana Alexander for initial help in scripting and Jodi Rowley for insightful discussions on various body size correction methods.

